# Quantitative live imaging reveals PRICKLE1 controls junctional neural tube morphogenesis independent of Planar Cell Polarity

**DOI:** 10.1101/2025.06.23.661208

**Authors:** Jian Xiong Wang, Yanina D. Alvarez, Siew Zhuan Tan, Samara N. Ranie, Samantha J. Stehbens, Melanie D. White

**Affiliations:** Institute for Molecular Bioscience, The University of Queensland, Australia; School of Biomedical Science, The University of Queensland, Australia; Australian Institute for Bioengineering and Nanotechnology, The University of Queensland, Australia

**Keywords:** Neurulation, Junctional neural tube, Neural tube defects, Live imaging, Planar Cell Polarity, PRICKLE1, Apical constriction, Morphogenesis, EMT, Avian embryo

## Abstract

Neurulation, the process that forms the neural tube—the precursor to the brain and spinal cord—is frequently disrupted in congenital malformations. Primary and secondary neurulation are integrated at a junctional zone, yet the cellular dynamics linking these programs remain unknown. Using high-resolution quantitative live imaging in transgenic quail embryos, we show that the junctional neural tube forms through two coordinated processes: mediolateral convergence and EMT-driven ingression of medial neuroepithelial cells. We demonstrate that PRICKLE1, a core PCP protein, orchestrates these behaviors independently of planar polarity cues. PK1 is enriched at the apical cortex of medial cells, where it drives actomyosin accumulation and apical constriction. This function is essential for both convergence and cell ingression but is uncoupled from classical PCP axis establishment. Our findings redefine the molecular basis of junctional neurulation and implicate impaired EMT as a central cause of localized neural tube defects.

## INTRODUCTION

Neurulation is a critical early embryonic process in higher vertebrates which forms the neural tube, the precursor to the brain and spinal cord^1^. Failures in neurulation cause severe congenital malformations called neural tube defects (NTDs) which are amongst the most common human birth defects. The neural tube forms from head-to-tail, through two fundamentally different morphogenetic processes^1^. The rostral neural tube forms through folding and fusing of the neuroepithelium in a process called primary neurulation^2^. However, the caudal neural tube forms differently through secondary neurulation. This is characterised by condensation of cells to form a rod-like structure which undergoes canalisation to generate a lumen^3^. It is critical to nervous system function that these differing morphogenetic processes are integrated to form a continuous neural tube. The spatiotemporal dynamics of this process have not been determined due to a lack of live imaging and species differences in neurulation.

A junctional transition zone where primary and secondary neurulation appear to occur simultaneously across the dorsoventral axis, has been identified in avian^4,5^ and human embryos^6,7^. More recently, a local morphogenetic process called junctional neurulation was proposed to occur in the avian junctional zone, ensuring continuity between the primary and secondary neural tubes^8^. Notably, most human spinal defects cluster near this junctional zone, highlighting its susceptibility to disruption^6^. In humans, impaired formation of the junctional neural tube leads to local spinal dysraphisms, recently named Junctional Neural Tube Defects (JNTDs)^9–12^. Patients with JNTDs exhibit structurally and functionally disconnected primary and secondary neural tubes, accompanied by lower limb deformities, impaired motor and sensory function and incontinence^9^.

Mouse embryos lack a junctional zone^3,13^, therefore our limited understanding of junctional neurulation has been derived from studies of avian embryos - primarily using fixed tissue sections or low resolution live imaging^8^. At the molecular level, the protein PRICKLE1 (PK1) has been implicated in junctional neurulation in avian embryos^8^. PK1 is a core protein in the Planar Cell Polarity (PCP) signalling pathway which controls convergent extension in vertebrates^14–16^. Planar-polarized enrichment of PCP proteins along mediolateral cell junctions drives actomyosin-dependent junction contraction and facilitates mediolateral cell convergence^17–20^. This PCP-dependent convergence of cells towards the embryonic midline is crucial to bend the neural plate and form the primary neural tube^19^. Disruption of other core PCP genes including *CELSR1-3*, *VANGL1/2* and *DVL2* can cause NTDs along the entire head-to-tail axis. *PRICKLE1* mutations, however, are predominantly associated with local spinal dysraphisms^21–25^. How PK1 disruption specifically affects the morphogenesis of the junctional neural tube remains unknown.

Here, we use high spatiotemporal resolution live imaging, targeted manipulations and quantitative cell tracking in transgenic quail embryos to characterise the real time cellular dynamics of junctional neurulation. We demonstrate that the junctional neural tube is formed by a combination of mediolateral cell convergence and epithelial-to-mesenchymal transition (EMT)-driven ingression of medial cells in the junctional zone. Disrupting PK1 impairs medial cell ingression and causes localized JNTDs by a novel PCP-independent mechanism that links EMT to neural tube closure. We find that PK1 localizes to the apical cortex of medial junctional zone cells, where it promotes actomyosin accumulation and apical constriction. Strikingly, PK1’s control of cortical F-actin accumulation is uncoupled from planar cell polarity, but essential for both mediolateral convergence and EMT-driven ingression. Our findings reformulate the molecular logic of junctional neurulation and show how failure of EMT can cause localized defects in the junctional neural tube.

## RESULTS

### Junctional neural tube closure requires PK1

To investigate the morphogenesis of the junctional neural tube, we used live imaging of a transgenic quail model^26,27^. The cellular material for the junctional zone between primary and secondary neurulation is generated in a region that is caudal and lateral to Hensen’s node and rostral to the primitive streak^28,29^, starting at the 5-somite stage (ss) in quail (Figure 1A, B). Live imaging of Lifeact-EGFP transgenic quail embryos^26^ reveals the junctional zone progressively converges from the 5ss until the 9-10ss when the posterior neuropore closes (Figure 1B, Video S1). The cells of the junctional zone are SOX2^+^ and give rise to both the junctional neural tube which forms between somites 18-27, and precursors for the secondary neural tube initiating from somite 28 (Figure 1C, D). To characterize the cellular dynamics driving junctional zone morphogenesis, we mosaically labelled cell nuclei by electroporating *TgT2(CAG:NLS-mCherry-IRES-GFP-CAAX)* quail embryos^27^ with a plasmid encoding H2B-iRFP670. High-resolution live imaging combined with nuclear tracking revealed that cells converge from the lateral region of the junctional zone (defined as the outermost 100 μm at each lateral edge), towards the medial region consisting of 100 μm on each side of the midline (927 cell tracks in 3 embryos, Figure 1E, F, Video S2). Within 3 hours, the average convergent displacement was 16.26 ± 0.46 μm per cell, however cells in the lateral junctional zone showed significantly more convergent displacement (19.66 ± 1.84 μm) than those in the medial region (7.50 ± 0.96 μm, Figure 1F). The cells in the lateral region also exhibited significantly stronger directionality towards the midline than those in the medial region (Figure 1G). Further analysis revealed a gradient of mediolateral convergence (Figure 1H). Immunofluorescent staining showed expression of PK1 at the 5ss which was significantly enriched in the junctional zone by the 9-10ss (Figure 1I). To disrupt PK1 expression, we electroporated a DNA construct encoding an intronic miRNA targeting *PRICKLE1* (or a scrambled control miRNA) upstream of an H2B-iRFP670 reporter^30^ into ∼1ss transgenic quail embryos expressing nuclear-mCherry and membrane-localised EGFP (*TgT2(CAG:NLS-mCherry-IRES-GFP-CAAX),* Figure S1A, B)^27^. We followed the embryos by live imaging for ∼15 hours, by which time the posterior neuropore of the control embryos was typically closed. Knocking down PK1 expression in the junctional zone disrupted junctional neurulation and prevented closure of the posterior neuropore (Figures 1J, K, S1C, Video S3). These results confirm that correct formation of the junctional neural tube relies on PK1.

**Figure 1.**
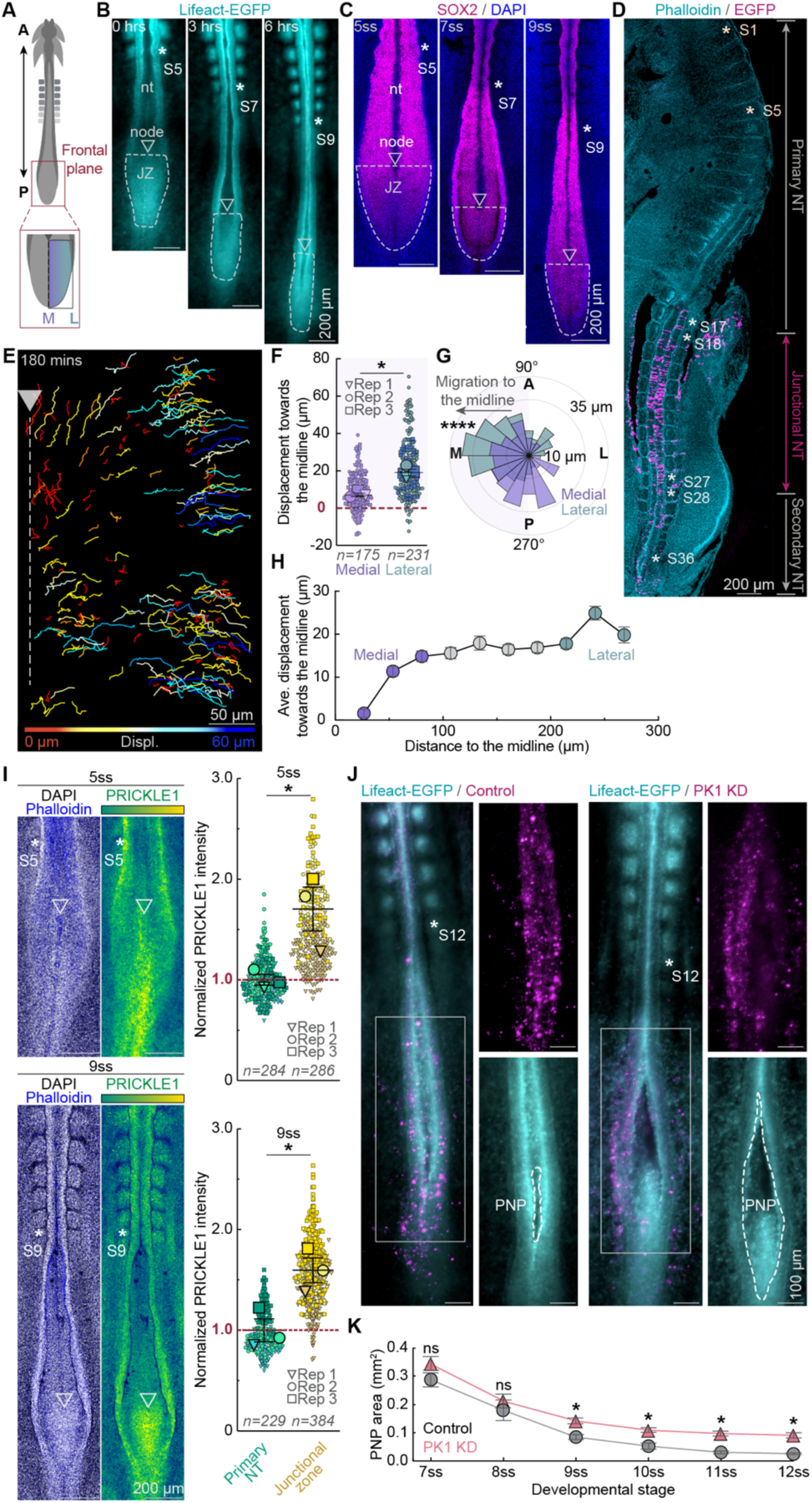
Junctional neural tube closure is PK1-dependent. (A) Schematic showing region of interest. (B) Location of junctional zone (JZ) from the 5ss to the 9ss in a Lifeact-EGFP embryo. (C) Location of JZ in fixed wildtype embryos stained with SOX2 and DAPI. (D) Wildtype embryo electroporated with EGFP mRNA in JZ and incubated for 48 hours. JZ cells contribute to junctional NT (somite 17/18 – 27/28) and secondary NT (>somite 27/28). (E) Snapshot showing individual tracks of migrating H2B-iRFP670 labelled cells. (F) Displacement towards the midline of cells in the medial and lateral regions. (G) Directionality of cell trajectories in the medial (n = 175 cells from 3 embryos) and lateral (n = 231 cells from 3 embryos) regions. ****p<0.0001, Kolmorogov-Sminorv test. (H) Cell displacement towards the midline plotted against cell starting positions across the mediolateral axis (927 tracked cells from 3 embryos, mean, ± sem, (I) PK1 is highly expressed in JZ compared to primary NT. (J, K) PK1 knockdown impairs posterior neuropore (PNP) closure. *p<0.05, un-paired t-test. F, I: * P<0.05, paired t-test. nt: neural tube. Asterisks indicate somite positions. See also Figure S1.

### PK1 knockdown disrupts cellular convergence without affecting planar cell polarity

To test how convergent cell movement within the junctional zone is impacted by PK1 knockdown, we electroporated ∼1ss *TgT2(CAG:NLS-mCherry-IRES-GFP-CAAX)* quail embryos with the plasmids encoding the PK1 miRNA or the scrambled control miRNA upstream of H2B-iRFP670. We performed live cell nuclei tracking from ∼5ss when junctional neurulation begins (Figure 2A, B). PK1 knockdown cells showed significantly less convergent displacement (Figure 2B-D, Video S4), and reduced directionality towards the midline (Figure 2E) compared to scrambled control cells.

**Figure 2.**
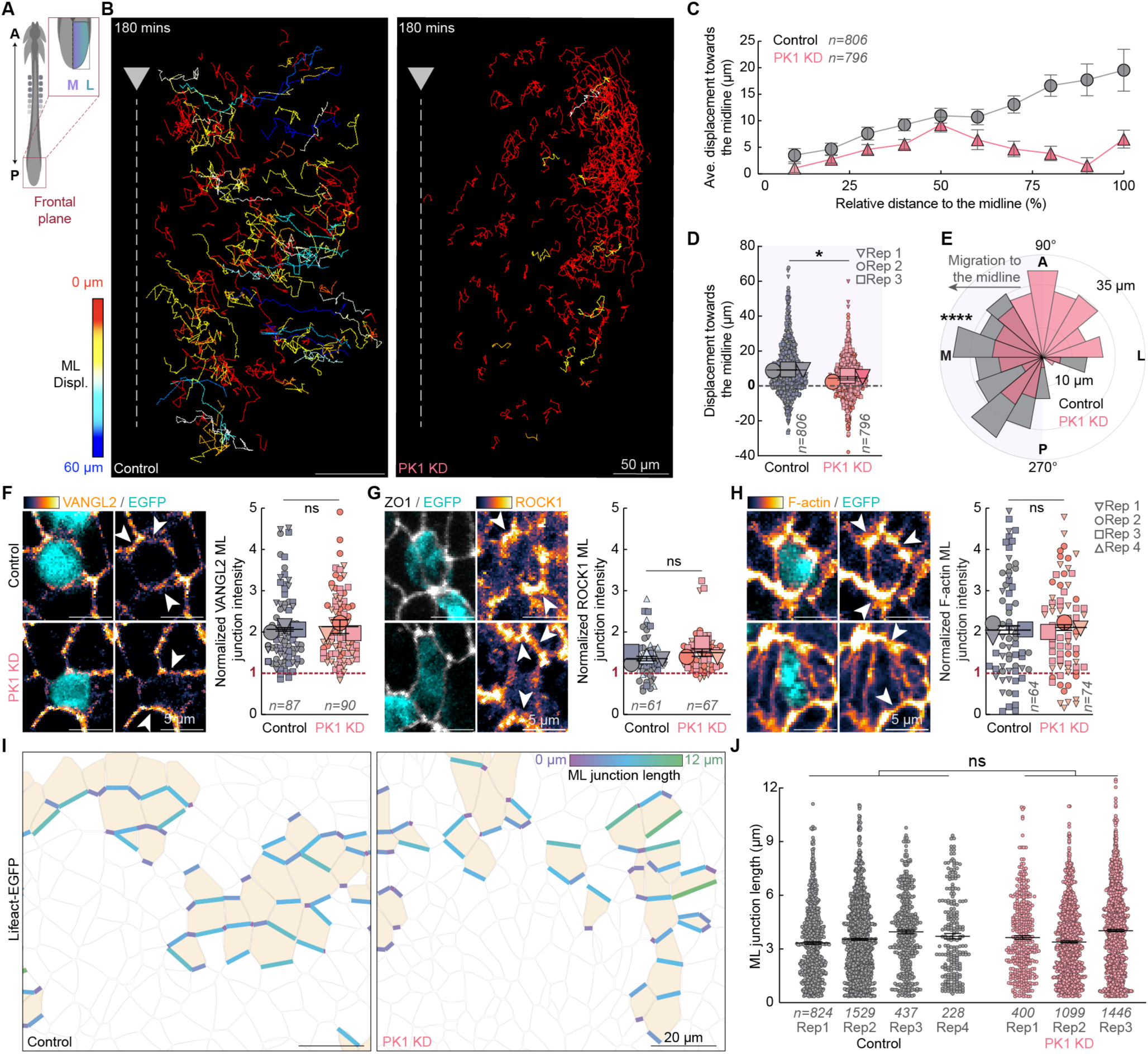
PK1 knockdown disrupts cellular convergence without affecting planar cell polarity. (A) Schematic shows region of interest. Snapshots show mediolateral trajectories of individual Scrambled control miRNA-H2B-iRFP670 or PK1 miRNA-H2B-iRFP670 labelled cells. (B) Cell displacement towards the midline plotted against cell starting positions across the mediolateral axis for PK1 knockdown (796 tracked cells from 3 embryos) and Control cells (n = 806 cells from 3 embryos, mean, ± sem). (C, D) Displacement towards the midline of PK1 knockdown (796 tracked cells from 3 embryos) and Control cells (n = 806 cells from 3 embryos), *p<0.05, un-paired t-test. (E) Directionality of PK1 knockdown and Control cells, ****p<0.0001, Kolmorogov-Sminorv test. (F-H) Immunofluorescent staining of PCP protein VANGL2 and downstream effectors ROCK1 and F-actin reveals no change in planar polarisation following PK1 knockdown. ML; mediolateral (I) Segmentation and identification of mediolaterally oriented junctions of PK1 knockdown or control cells in the JZ. (J) PK1 knockdown does not alter mediolateral junction length, ns, not significant, unpaired t-test. See also Figure S2.

As PK1 is a core PCP protein, we determined whether the reduction in cell convergence following PK1 knockdown results from disrupted planar cell polarity. Immunofluorescent staining confirmed the PCP components VANGL2 and PK1 and their downstream effectors, ROCK1, pMLC and F-actin, are planar polarized, characterized by enrichment along mediolateral cell junctions in the junctional zone of the ∼5ss quail embryo (Figure S2A-C). However, unexpectedly, PK1 knockdown did not alter the planar polarity of VANGL2, ROCK1 or F-actin (Figure 2F-H). Furthermore, PK1 knockdown had no effect on the length of mediolateral cell junctions in the junctional zone (Figures 2I, J, S2D-F), indicating PCP signalling remained intact. Together, these results suggest PK1 has a PCP-independent role in junctional neural tube formation.

### PK1 knockdown disrupts medial cell ingression

To understand how PK1 knockdown disrupts cell convergence without affecting planar cell polarity in the junctional zone, we examined cellular movements in the medial region, where cells typically converge to. By analysing the dorsoventral displacement of the H2B-iRFP670 labelled cell nuclei tracked in Figure 1E, we found that cells within the medial region ingressed from the dorsal surface of the junctional zone into the underlying ventral tissue (Figure 3A, B, Video S5). Like the convergent displacement behavior, we observed a gradient of ventral displacement: within 3 hours of live imaging, cells in the medial region moved ventrally (6.30 ± 0.70 μm), while cells in the lateral region moved dorsally (3.49 ± 1.02 μm, Figure 3C, D). The medial cells also exhibited significantly stronger ventral directionality than cells in the lateral regions (Figure 3E). Live imaging of the dorsal surface of the junctional zone in Lifeact-EGFP quail embryos at ∼5ss enabled visualization of the apical cell surfaces (Figure 3F). Cells in the medial region accumulated cortical LifeAct-EGFP and reduced their apical surface area over time before ingressing and disappearing from the dorsal surface (Figures 3F,G). Significantly more cells ingressed in the medial region than the lateral regions (Figure 3H).

**Figure 3.**
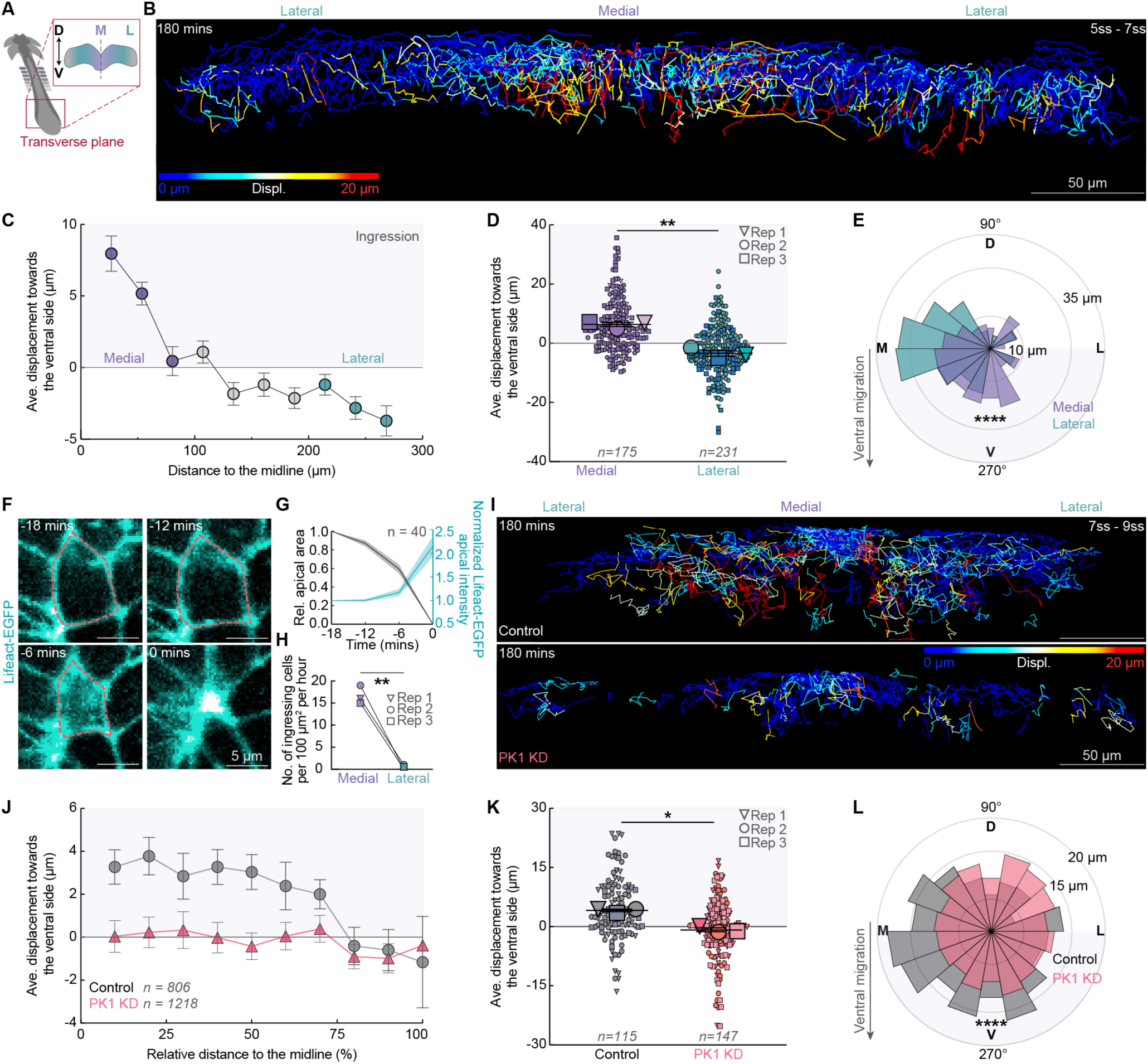
PK1 knockdown disrupts medial cell ingression. (A) Schematic showing region of interest. (B) Snapshot showing individual tracks of H2B-iRFP670 labelled cells migrating along the dorsoventral axis. (C) Ventral cell displacement plotted against cell starting positions across the mediolateral axis (927 tracked cells from 3 embryos, mean, ± sem). (D) Ventral displacement of cells in the medial (n = 175 cells from 3 embryos) and lateral (n = 231 cells from 3 embryos) regions. **p<0.01, paired t-test. (E) Directionality of cell trajectories along the dorsoventral (D, V) and mediolateral (M, L) axes in the medial (n = 175 cells from 3 embryos) and lateral (n = 231 cells from 3 embryos) regions. ****p<0.0001, Kolmorogov-Sminorv test. (F) Accumulation of Lifeact-EGFP at the shrinking apical cortex of an ingressing cell. (G) Lifeact-EGFP accumulates at the apical cortex as the apical cell area shrinks. (H) Significantly more cells ingress in the medial JZ region than the lateral, n = 3 embryos, *p<0.05, paired t-test. (I) Snapshots showing dorsoventral trajectories of individual Scrambled control miRNA-H2B-iRFP670 or PK1 miRNA-H2B-iRFP670 labelled cells. (J) Ventral displacement plotted against cell starting positions across the mediolateral axis for PK1 knockdown (1218 tracked cells from 3 embryos) and Control cells (n = 806 cells from 3 embryos, mean, ± sem). (K) Ventral displacement of PK1 knockdown and Control cells, *p<0.05, un-paired t-test. (L) Directionality of PK1 knockdown (n = 115 cells from 3 embryos) and Control cells (n = 147 cells from 3 embryos) along the dorsoventral (D, V) and mediolateral (M, L) axes, ****p<0.0001, Kolmorogov-Sminorv test.

To determine if PK1 knockdown affects the medial cell ingression, we electroporated ∼1ss embryos with the PK1 miRNA-H2B-iRFP670 construct (or the scrambled control) and tracked cell nuclei by live imaging from ∼5ss. PK1 knockdown significantly reduced cell ingression in the medial region compared to scrambled controls (Figure 3I-K, Video S6). This was accompanied by a significant loss of ventral directionality (Figure 3L), confirming that PK1 is required for cell ingression in the medial region of the junctional zone.

### Medial cells undergo an epithelial-to-mesenchyme transition

PK1 has not been previously linked to cell ingression, therefore we conducted a more detailed investigation into the mechanisms causing medial cells to ingress. The junctional zone is situated immediately rostral to the primitive streak: a region of ongoing cell ingression. Cells in the primitive streak undergo an epithelial-to-mesenchymal transition (EMT) during which they constrict their apical surfaces and ingress into the deeper tissue layer^31,32^. Therefore, we hypothesized that a similar mechanism may drive the ingression of cells in the medial region of the junctional zone.

The *SNAIL* gene family member, SLUG, is essential for EMT and cell ingression at the primitive streak^33^. Immunofluorescent staining revealed SLUG^+^ cells localized to the medial region of the junctional zone (Figure 4A). To confirm that the SLUG^+^ cells in the medial region ingress, we used a SLUG enhancer construct that reports SLUG expression^34^ (Figure S3A). We electroporated 1ss Lifeact-EGFP quail embryos with a DNA plasmid that expresses membrane-mScarlet under the control of the SLUG enhancer (SLUG enh-memb-mScarlet) and followed the cells by live imaging from ∼5ss. Many cells within the junctional zone were labelled with SLUG enh-memb-mScarlet and visibly reduced their apical surface area over time before disappearing from the dorsal side (Figure 4B). The SLUG enhancer-positive cells were initially distributed across the junctional zone but within 6 hours, moved to within 50 μm of the midline, where they ingressed (Figure 4C, D, Video S7). This medial movement was significantly faster than cells labelled with a ubiquitous CMV promoter (Figure S3B). Furthermore, there was a morphological change from an epithelial profile in the lateral region to a highly protrusive mesenchymal morphology in the medial region (Figures 4E, F, S3C).

**Figure 4.**
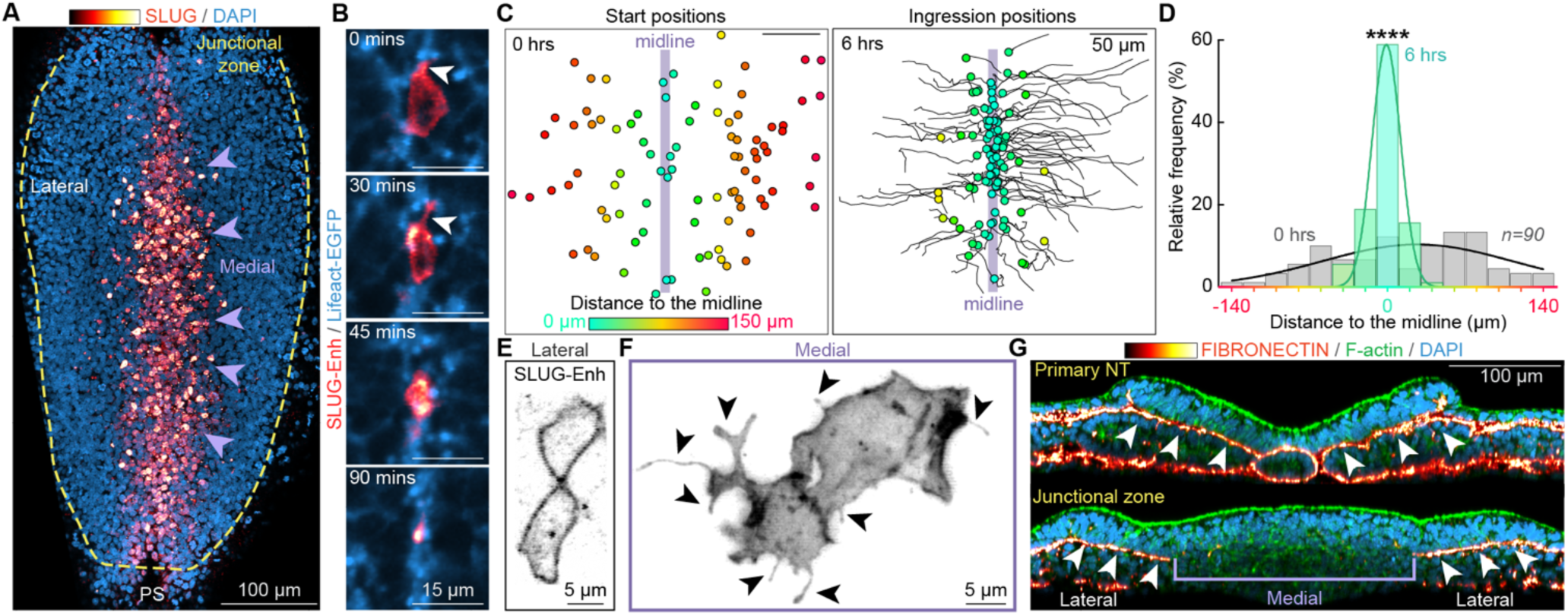
Medial junctional zone cells undergo EMT and ingress. (A) Immunofluorescent staining showing SLUG expression in medial JZ cells (arrowheads). PS; primitive streak. (B) Live imaging of a medial cell expressing a SLUG reporter constricting its apical surface, extending protrusions (arrowheads) and ingressing. (C) Positions of SLUG enhancer-positive cells at the onset of live imaging (0 hours, left) and their ingression sites near the midline (right). SLUG enhancer-positive cells are initially distributed across the medial region of the JZ (grey bars), but ingress near the midline (green bars), (n = 90 cells from 4 embryos). ****p<0.0001, Kolmorogov-Sminorv test. (E) SLUG enhancer-positive cell morphology is epithelial in the lateral JZ. (F) SLUG-enhancer positive cells have a highly protrusive (arrowheads) mesenchymal morphology in the medial JZ. (G) Immunofluorescent staining reveals a continuous layer of FIBRONECTIN underneath the primary neural plate (top, arrowheads) which is absent in the medial region of the JZ (bottom, bracket). See also Figure S3.

Another crucial early step in EMT in the primitive streak is degradation of the basement membrane^31^. During primary neurulation, the ventral side of the neural plate is covered by a layer of the extracellular matrix protein FIBRONECTIN (Figure 4G, top). This FIBRONECTIN layer remains intact in the lateral regions of the junctional zone but is absent in the medial region, consistent with EMT-driven medial cell ingression (Figure 4G, bottom). Together, these findings demonstrate that cells in the junctional zone of the neural plate undergo an EMT once they reach the medial region to ingress towards the ventral side.

### Medial cell ingression is required for junctional neural tube formation

We next asked whether the medial cell ingression is necessary for the correct formation of the junctional neural tube. To block EMT and medial cell ingression, we co-electroporated a morpholino targeting SLUG (or a non-targeting control morpholino) with a plasmid encoding H2B-EGFP into the junctional zone at the 1ss. We then fixed and stained the embryos for SLUG and DAPI during junctional neurulation at the 7ss (Figure S4A). In control embryos, 37.2 ± 7.2 % (mean ± s.d.) of the electroporated cells in the medial junctional zone were SLUG^+^. This was reduced to 8.6 ± 1.3 % of electroporated medial cells in the SLUG knockdown embryos, confirming the efficacy of the morpholino (Figure 5A). In control embryos, the H2B-EGFP-positive cells were distributed across the dorsoventral axis of the junctional zone and 23.6 ± 6 % of cells ingressed to the ventral third of the tissue (Figure 5B, C). However, in SLUG knockdown embryos, the electroporated cells remained significantly closer to the dorsal surface and only 11.9 ± 5 % of H2B-GFP-positive cells penetrated to the ventral third of the junctional zone (Figure 5B, C). These results confirm that SLUG expression is required for medial cell ingression.

**Figure 5.**
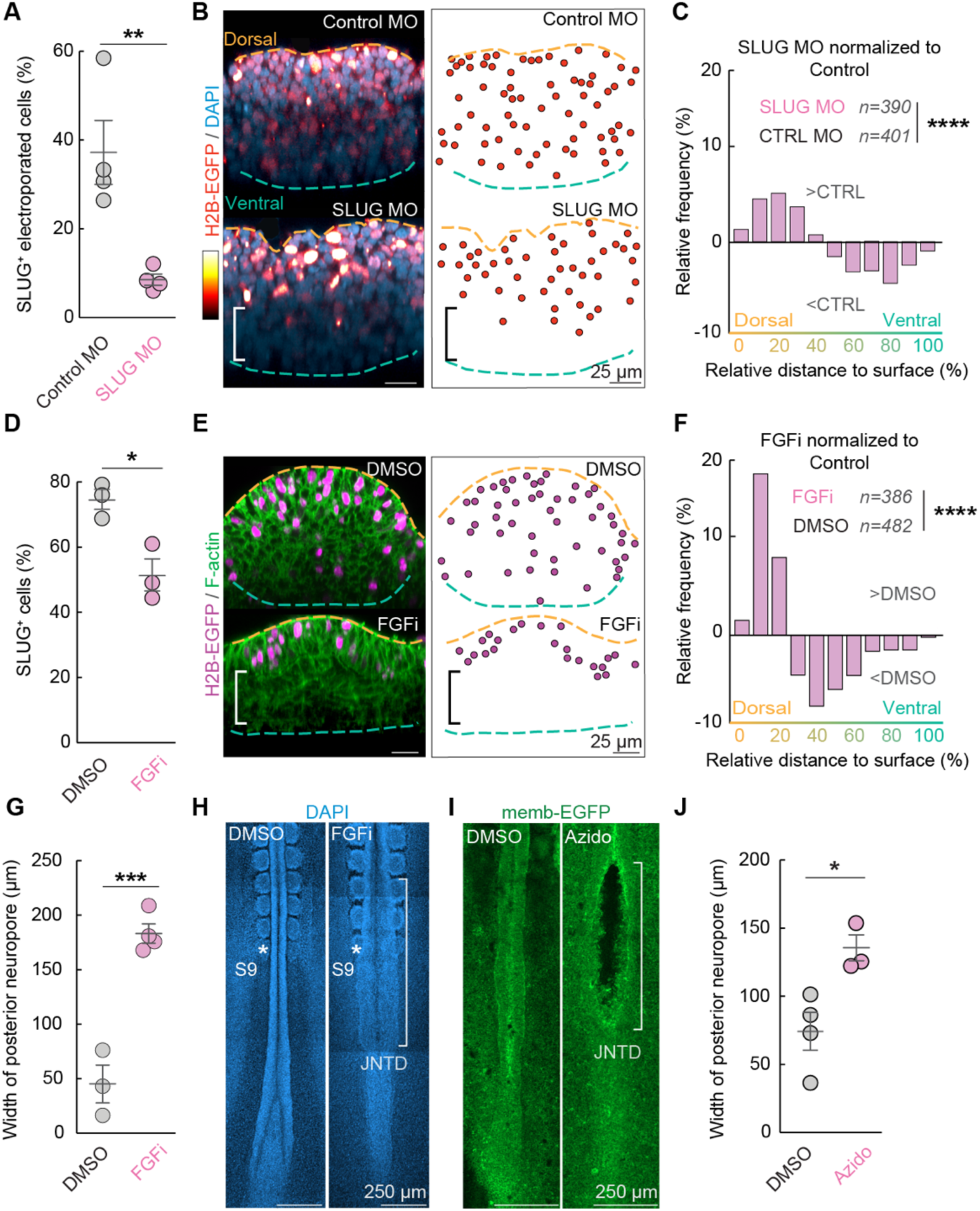
Medial cell ingression is required for junctional neural tube formation. (A) Electroporation of a SLUG morpholino (MO) effectively reduces the number of SLUG^+^ cells in the JZ. **p<0.01, unpaired t-test. (B) SLUG knockdown reduces ingression of electroporated cells to the ventral side (bracket). (C) Relative frequencies of SLUG knockdown cells normalized to control cells along the dorsoventral axis. SLUG knockdown cells (390 cells from 4 embryos) are significantly more frequently located near the dorsal surface than Control cells (401 cells from 4 embryos). ****p<0.0001, Kolmorogov-Sminorv test. (D) FGF inhibition significantly reduces the number of SLUG^+^ cells in the JZ. * p<0.05, unpaired t-test. (E) FGF inhibition prevents ingression of electroporated medial JZ cells (magenta) to the ventral side of the tissue (bracket). (F) Relative frequencies of cells in embryos treated with FGF inhibition normalized to DMSO control along the dorsoventral axis. Cells in FGF inhibitor-treated embryos (386 cells from 3 embryos) are significantly more frequently located near the dorsal surface than DMSO control-treated cells (482 cells from 3 embryos). ****p<0.0001, Kolmorogov-Sminorv test. (G) The posterior neuropore is significantly wider in 9 – 10 somite stage FGF inhibitor-treated embryos (n = 4) than the DMSO-treated controls (n = 3). ***p<0.001, unpaired t-test. (H) FGF inhibition causes a junctional neural tube defect (JNTD, white bracket). (I) Photoactivating azidoblebbistatin in the medial JZ causes a JNTD (white bracket). (J) Photoactivating azidoblebbistatin in the medial JZ results in a significantly wider posterior neuropore (n = 3 embryos) compared to DMSO-treated controls (n = 4 embryos). See also Figure S4.

As electroporation results in mosaic expression, it is challenging to reliably target the entire subset of SLUG^+^ cells in the junctional zone. Therefore, we turned to pharmacological inhibition to block medial cell ingression. Fibroblast Growth Factor (FGF) signalling is required for the expression of SLUG and cell ingression at the primitive streak in mouse and chick embryos^35,36^. We treated 1 ss wildtype embryos with 10 mM of the FGF Receptor inhibitor Infigratinib^37^ or a DMSO control and then electroporated the junctional zone with the H2B-EGFP plasmid to assay for cell ingression. To confirm that blocking FGF signalling reduced SLUG expression, we fixed embryos at the 5ss when junctional neurulation initiates and stained for SLUG. In the control embryos, 74.7 ± 5.3% (mean ± s.d.) of cells in the medial junctional zone were SLUG+, whereas this reduced to 51.5 ± 8.5 % in the FGF-inhibited embryos (Figure 5D). Consistent with the effects of reducing SLUG expression, in the FGF-inhibited embryos the H2B-EGFP labelled cells remained significantly closer to the dorsal surface of the junctional zone than in the DMSO control embryos and no cells penetrated to the ventral third of the tissue (Figure 5E, F). Furthermore, by the 9-10ss the posterior neuropore had typically closed in control embryos with an average width of 45.40 ± 17.27 μm. By contrast, the posterior neuropore remained open in FGF-inhibited embryos (average width of 183.5 ± 8.912 μm), resulting in a junctional neural tube defect (Figure 5G, H). Blocking FGF signalling did not affect the assembly of actin cables associated with convergence in the lateral regions of the junctional zone (Figure S4B), suggesting the defect primarily results from the prevention of medial cell ingression.

Prior to ingression, the medial cells undergo apical constriction (Figure 3F) – a process that is dependent on actomyosin contraction^38^. Therefore, to target cell ingression in the medial region of the junctional zone only, we treated *TgT2(CAG:NLS-mCherry-IRES-GFP-CAAX)* embryos at the 5ss with a photo-activatable myosin II inhibitor, azidoblebbistatin^39,40^. We specifically activated the azidoblebbistatin in the medial region using a 405 nm laser. Consistent with the FGF inhibition, blocking actomyosin contraction in only the medial cells caused focalised junctional NTDs, characterised by a significantly wider posterior neuropore at the ∼16ss (Figure 5I, J). Thus, medial cell ingression is required for the correct formation of the junctional neural tube.

### PK1 regulates cortical actomyosin in a PCP-independent manner for medial cell ingression

To understand PK1’s role in medial cell ingression we first tested whether PK1 controls SLUG expression. We electroporated 1ss wildtype quail embryos with the plasmids encoding the intronic PK1 miRNA or the scrambled control miRNA upstream of EGFP. We fixed and immunostained the embryos for SLUG at 5ss and found that knocking down PK1 did not change SLUG levels. This confirms that PK1 does not regulate SLUG expression in the junctional zone (Figure S5A,B).

To investigate how PK1 knockdown affects the cell apical surface area, we expressed the PK1 miRNA or the scrambled control miRNA upstream of membrane-mScarlet in the junctional zone of Lifeact-EGFP quail embryos and examined the distribution of apical cell surface size at the 5ss. We used a custom local Z projection strategy to project the curved dorsal surface of the junctional zone onto a 2D plane and computationally segmented cell boundaries labelled by Lifeact-EGFP^26^. Quantification revealed the average apical surface area of PK1 knockdown cells was significantly larger than scrambled control cells (Figure 6A, B). To determine if the dilated apical cell area results from defects in apical constriction, individual cells were tracked for ∼3 hours and the change in apical surface area over time was analysed^41^. A closer examination of the medial cell apical area dynamics revealed no differences between control cells and PK1 knockdown cells during the non-constriction phase. However, during the constriction phase, PK1 knockdown cells displayed a significantly smaller reduction in apical surface area compared to controls (Figure 6C-E, Video S8). In the control embryos, all tracked medial cells shrunk their apical surfaces within 3 hours and 40% of them fully ingressed (Figure 6C, D). By contrast, in the PK1 knockdown embryos, 90% of the tracked medial cells reduced their apical surface by less than 35%, and none shrank by more than 55% (Figure 6C-E). Furthermore, none of the PK1 knockdown cells ingressed within the 3 hour observation period. These findings demonstrate that PK1 knockdown specifically inhibits medial cell apical constriction, preventing ingression.

**Figure 6.**
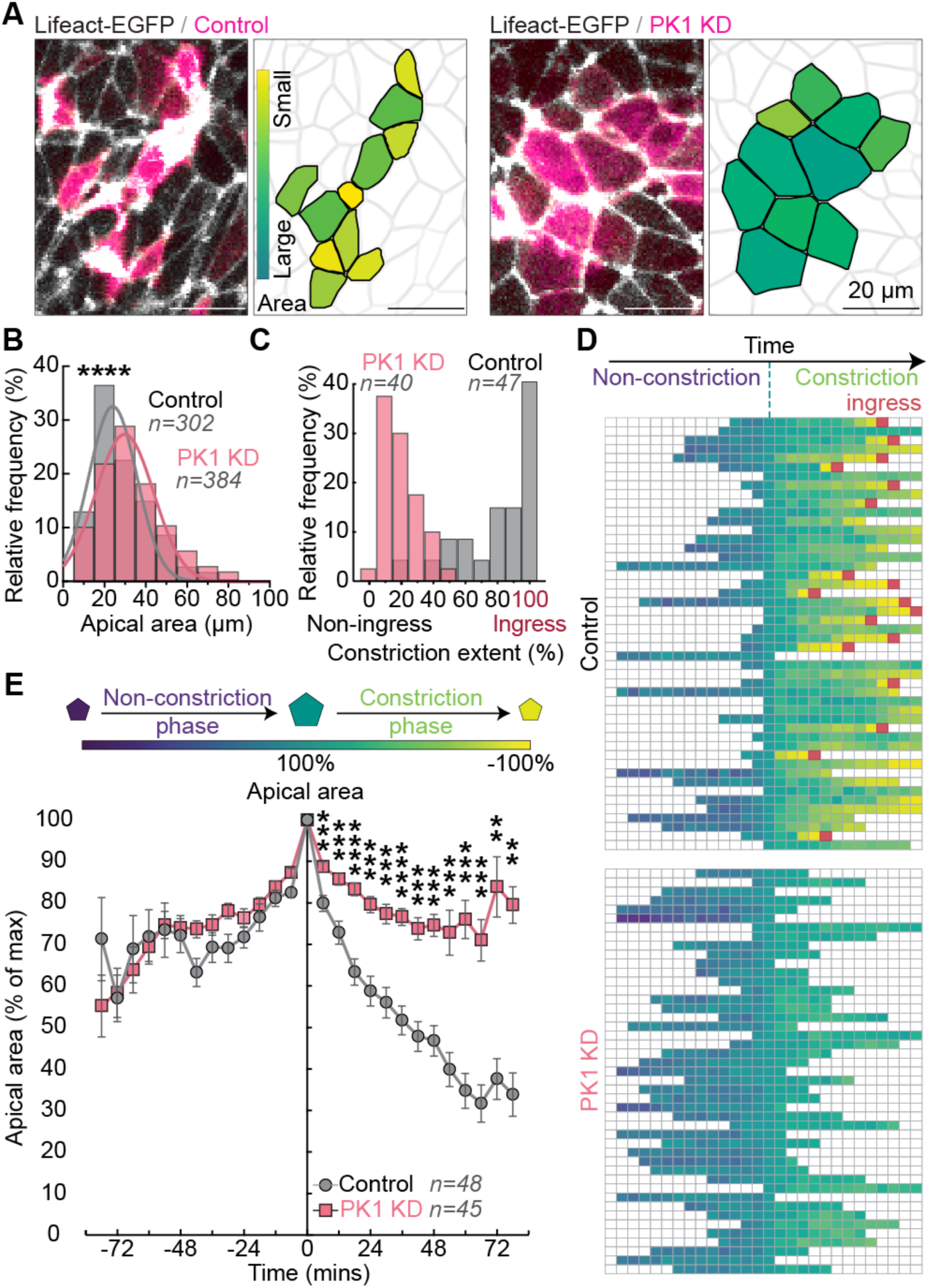
Ingression of medial junctional zone cells requires PK1-dependent apical constriction. (A, B) PK1 knockdown causes a significant increase in JZ cell apical area. (n = 384 cells from 3 PK1 knockdown embryos and n = 302 cells from 3 control embryos). ****p<0.0001, Kolmorogov-Sminorv test. (C) PK1 KD cells (n = 40 cells from 3 embryos) constrict their apical surface less than Control cells (n = 47 cells from 3 embryos) and fail to ingress. (D) Individual Control and PK1 KD cells colour-coded by apical surface area over time, aligned by their maximal apical area. The red squares indicate ingression events. (E) Quantification of cell apical area during live imaging. Individual cells are aligned to t=0 mins when they reach their maximal area. The negative time refers to the preceding non-constriction phase, and the positive time represents the constriction phase when the cell apical surface shrinks (n=45 cells from 3 PK1 knockdown embryos and 48 cells from 3 control embryos), **p<0.01, ***p<0.001, ****p<0.0001, unpaired t-test. See also Figure S6.

Apical constriction is associated with enrichment of actomyosin at the apical cortex^38^ and PK1 has recently been shown to regulate cortical F-actin in *Xenopus* embryos^42^. To characterise actomyosin at the apical cortex in the junctional zone, we fixed wildtype quail embryos at the 5ss and immunostained for activated Myosin Light Chain (p-MLC), F-actin and N-CADHERIN to label the cell junctions (Figure 7A, B). Large patches of p-MLC and F-actin were present at the apical cortex of 22 – 69% of medial cells (Figure 7C). Although the proportion of medial cells with apical actomyosin patches varied among embryos, nonetheless, there were always significantly fewer cells with enriched apical actomyosin in the lateral regions of the junctional zone (2.5 – 10.5% of lateral cells, 4 embryos, Figure 7C). In *Xenopus* mesodermal cells, it is the diffuse cytoplasmic pool of PK1 that increases cortical F-actin^42^. Therefore, we examined PK1 subcellular localisation near the cortex of cells in the junctional zone. Immunofluorescent staining showed significantly higher levels of PK1 at the apical cortex of medial cells compared to lateral cells in the junctional zone (Figure 7D – G). Enrichment of apical F-actin was often observed in the medial cells with high cortical PK1 (Figure 6F). To understand how PK1 affects the cortical accumulation of actin in the quail embryo in real time, we electroporated the junctional zone of Lifeact-EGFP embryos with the PK1 miRNA or the scrambled control miRNA upstream of membrane-mScarlet at the 1ss and performed live imaging from 5ss. In the scrambled control cells, there was a significant increase in Lifeact-EGFP intensity at the apical cortex, concomitant with a significant reduction in apical surface area (Figures 7H, I, S6). The PK1 knockdown cells, however, did not show any change in Lifeact-EGFP intensity or apical surface area, confirming that PK1 is required for cortical accumulation of Lifeact-EGFP and apical constriction.

**Figure 7.**
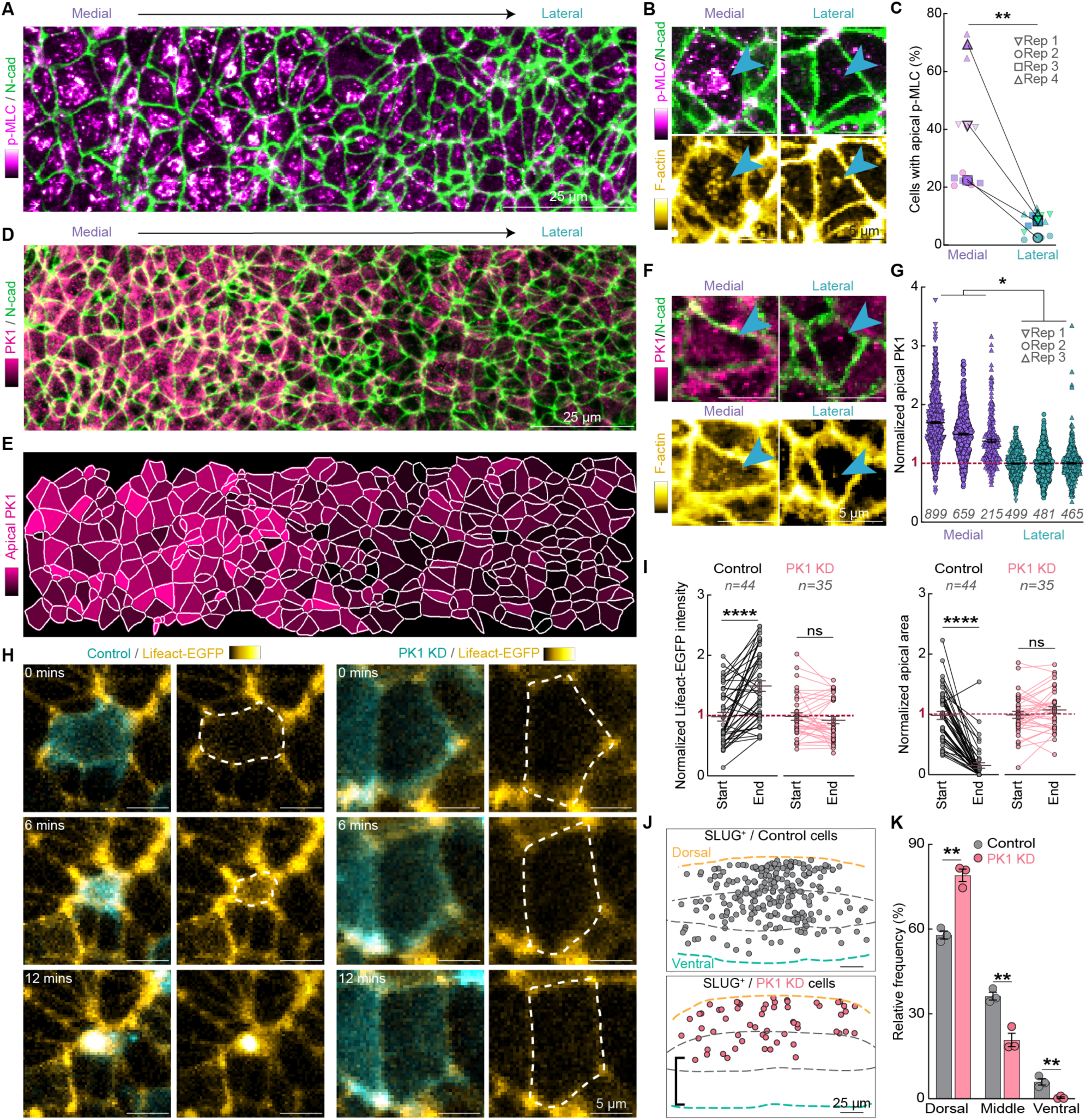
PK1 regulates cortical actomyosin for apical constriction. (A) Immunofluorescent staining showing the differential distribution of apical p-MLC (magenta) across the mediolateral axis of the JZ. (B) Apically enriched p-MLC is observed in medial but not in lateral cells (cyan arrow heads, upper panel). The cell with apically enriched p-MLC in the medial region also shows enrichment of apical F-actin (cyan arrowheads, lower panel). (C) Comparison of the percentage of cells with apical p-MLC accumulation between medial and lateral region (n = 4 embryos), **p<0.01, paired t-test. (D) Immunofluorescent staining showing PK1 localisation at the apical cortex of cells in the JZ. (E) Heatmap of PK1 intensity at the apical cortex in the JZ. (F, G) Medial cells have significantly more PK1 at the apical cortex than lateral cells (blue arrowheads, n = 3 embryos). *p<0.05, paired t-test. (H) Live imaging shows Lifeact-EGFP accumulates at the apical cortex of a Control cell undergoing ingression but not in a PK1 KD cell. (I) Quantification showing Lifeact-EGFP significantly increases at the apical cortex of Control cells (n = 44 cells from 3 embryos) as they shrink their apical area. PK1 KD cells (n = 35 cells from 3 embryos) show no change in Lifeact-EGFP intensity or apical area. ****p<0.0001, ns: not significant, paired t-test. (J,K) PK1 KD results in a significantly higher frequency of SLUG^+^ cells in the dorsal third of the JZ and significantly fewer SLUG^+^ cells in the middle and ventral thirds of the JZ compared to Control SLUG^+^ cells. **p<0.01, unpaired t-test. See also Figure S6.

Our findings reveal that correct formation of the junctional neural tube requires 1) ingression of SLUG^+^ medial cells and 2) PK1-dependent cortical actin accumulation and apical constriction. However, PK1 does not alter SLUG expression. This led us to finally ask if SLUG^+^ cells rely on PK1-mediated apical constriction to ingress. To address this question, we electroporated wildtype embryos with the PK1 miRNA or the scrambled control miRNA upstream of H2B-iRFP670 at the 1ss. We then fixed and immunostained for SLUG at the 7ss to identify ingressing cells. Quantitative analysis demonstrated that whereas SLUG^+^/Control cells were distributed throughout the dorsoventral axis of the junctional zone, SLUG^+^/PK1 KD cells remained significantly closer to the dorsal surface and did not penetrate at all to the most ventral tissue (Figure 7J, K). These results demonstrate that SLUG^+^ medial cells rely on PK1-driven apical constriction to successfully ingress. Disruption of PK1 impairs SLUG^+^ cell apical constriction and ingression, ultimately leading to junctional neural tube defects.

## DISCUSSION

Junctional neural tube closure is a critical morphogenetic event bridging primary and secondary neurulation. By combining high resolution live imaging, targeted manipulations, and quantitative cell tracking in transgenic quail embryos, we reveal that PK1 is necessary for both robust mediolateral cell convergence and for the apical constriction–dependent ingression of medial junctional zone cells. Importantly, these functions of PK1 are executed independently of classical PCP signalling. Therefore, we have established a novel, PCP-independent role for PK1 in neural tube morphogenesis. Given that impaired junctional neural tube formation underlies human junctional neural tube defects^9–12^, and that *PRICKLE1* mutations have been linked to local spinal dysraphisms^21^, our findings provide a mechanistic basis for how PK1 dysfunction may contribute to human disease.

### Medial cell ingression couples EMT to neural tube closure

Our live imaging and cell tracking reveal that junctional zone cells initially converge mediolaterally before undergoing dorsoventral ingression into the underlying ventral tissue (Figures 2, 3). Medial cells express the EMT regulator SLUG, constrict their apical surfaces and adopt protrusive, motile morphologies reminiscent of cells ingressing at the primitive streak (Figure 4). Disruption of SLUG function, either by morpholino knockdown or FGF-inhibition, prevents medial cell ingression and yields focal junctional neural tube defects (Figure 5), demonstrating that EMT-like behaviour is essential for neural tube closure. Traditionally, EMT-mediated ingression at the primitive streak is viewed as a hallmark of gastrulation, distinguishing mesendodermal-fated cells from adjacent non-ingressing neuroectoderm^43^. However, in the junctional zone we found that medial neuroectoderm cells also undergo a similar EMT and ingress. Yet, these cells largely retain neural identity and ultimately contribute to both the junctional and secondary neural tubes (Figure 1D). This uncoupling of EMT from lineage transition suggests that, unlike in the primitive streak, EMT within the neural plate primarily serves a morphogenetic function—facilitating tissue remodelling and neural tube elongation—rather than driving a lineage transition.

### A novel, PCP-independent role for PK1

PK1 is traditionally classified as a core component of the PCP pathway, where it contributes to convergent extension movements in multiple vertebrate systems^44–47^. However, our data demonstrate that PK1 knockdown does not disrupt the planar-polarized distribution of VANGL2, ROCK1, or F-actin at mediolateral junctions in the junctional zone, nor does it affect junction length (Figure 2). Instead, PK1 is specifically enriched at the apical cortex of medial junctional zone cells, where it is essential for the localized accumulation of F-actin and activated myosin - key drivers of apical constriction (Figure 7). This functional uncoupling of PK1 from canonical PCP axis establishment reveals that, within the junctional zone, PK1 drives morphogenetic cell behaviors through a distinct mechanism. Our high resolution imaging confirms that apical F-actin accumulation is PK1-dependent in medial junctional zone cells (Figure 7), suggesting that PK1 directly modulates cortical actin dynamics, as recently described in *Xenopus*^42^. Given that FGF signalling does not regulate PK1 expression in avian embryos^48^, we propose that PK1 functions in parallel with the EMT program, coupling actomyosin–driven apical constriction to neuroepithelial cell ingression.

PK1 is broadly expressed throughout vertebrate embryogenesis and studies in mouse, *Xenopus*, zebrafish and avian embryos have demonstrated its requirement in neurulation and the development of cardiac, respiratory, renal, skeletal, dental, auditory, and ocular structures^49^. Disruptions in these processes resulting from PK1 perturbations are typically attributed to PK1’s function in the PCP pathway. However, our findings, together with growing evidence of PCP protein functions independent of planar polarity^42,50,51^, suggest that PK1 may also regulate the actomyosin cortex and promote apical constriction—a key morphogenetic process—in contexts beyond junctional neurulation.

The PCP-independent role of PK1 in the junctional zone may explain why PK1 disruptions lead to localized NTDs specifically at the site of neuroepithelial cell ingression, in contrast to mutations in other core PCP proteins such as CELSR1-3 and VANGL1/2 which can impair neural tube closure along the anteroposterior axis^22–25^. We propose that the primary defect following PK1 disruption in the junctional zone arises from impaired medial cell ingression. This is supported by our finding that targeted inhibition of actomyosin contractility in the medial region alone is sufficient to induce JNTDs (Figure 5). Therefore, the observed defects in lateral convergence may represent a secondary consequence of the failure to clear medial cells. Together, our findings not only advance our understanding of the cellular choreography underlying junctional neurulation but also provide mechanistic insight into the etiology of human junctional neural tube defects.

## Supporting information

Video S1. JZ in Lifeact-EGFP quail embryo converges

Video S2. Cells in JZ converge

Video S3. PK1 knockdown abolishes PNP closure

Video S4. PK1 knockdown disrupts cellular convergence

Video S5. Cells in medial JZ ingress

Video S6. PK1 KD damages medial cellular ingression

Video S7. SLUG positive cells migrate medially and ingress

Video S8. PK1 knockdown perturbs cell apical constriction and ingression

## ACKNOWLEDGEMENTS

Microscopy was performed at the Institute for Molecular Bioscience Microscopy Facility which was established with the support of the Australian Cancer Research Foundation (ACRF) and incorporates the Dynamic Imaging, Cancer Biology Imaging and Cancer Ultrastructure and Function Facilities. We thank the IMB Microscopy facility staff. M.D.W was supported by a Future Fellowship (FT200100899) and a Discovery Project grant (DP220101878) from the Australian Research Council (ARC), and Ideas Grants (2013027, 2038843) from the National Health and Medical Research Council of Australia (NHMRC).

## AUTHOR CONTRIBUTIONS

**JXW**; investigation, formal analysis, methodology, software, data curation, visualization, writing – review and editing. **YDA**; formal analysis, methodology, software, data curation, writing – review and editing. **SZT**; methodology, writing – review and editing. **SNR**; investigation. **SJS**; writing – review and editing. **MDW**; conceptualization, data curation, formal analysis, funding acquisition, methodology, project administration, supervision, validation, visualization, writing – original draft preparation, writing – review and editing.

## DECLARATION OF INTERESTS

The authors declare no competing interests.

## SUPPLEMENTAL FIGURES

**Figure S1.**
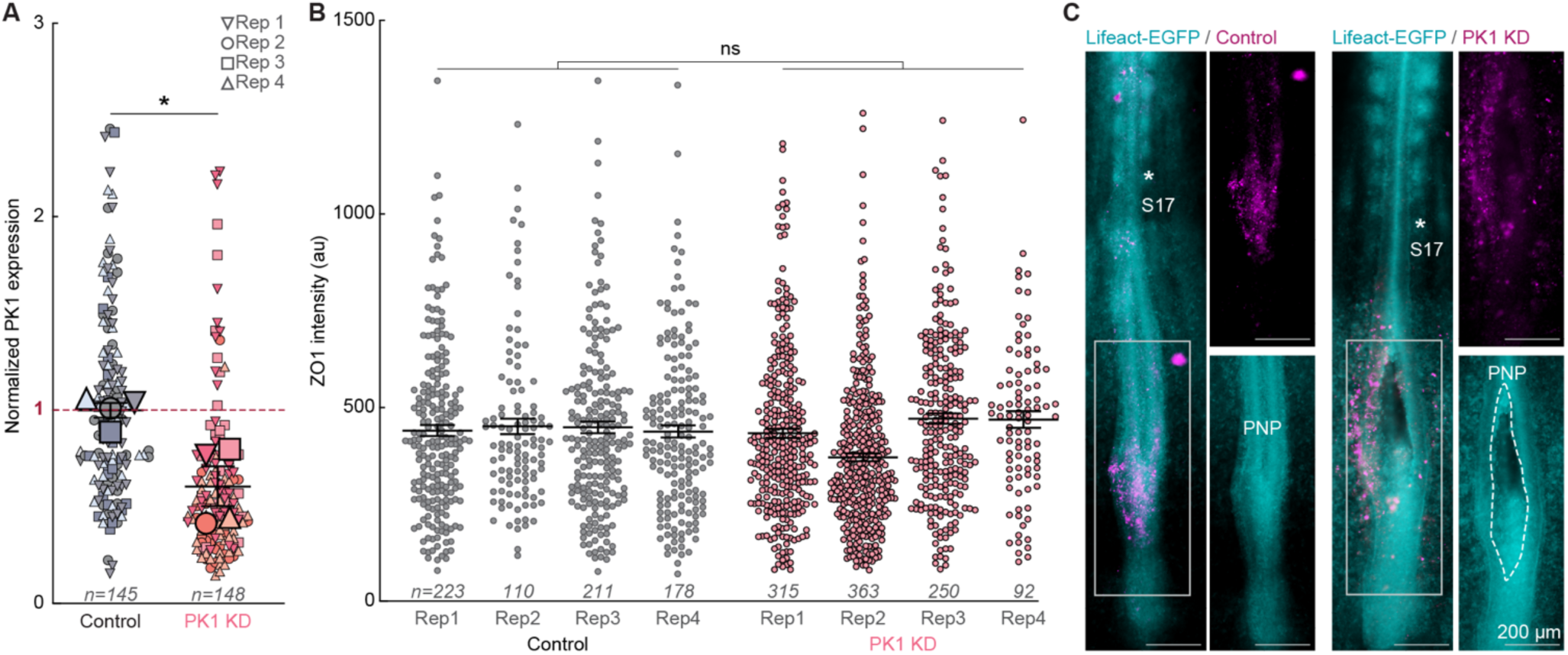
PK1 knockdown causes sustained disruption of neural tube closure, related to Figure 1. (A) PK1-knockdown miRNA decreases the endogenous PK1 expression level in the JZ. The expression level of PK1 is normalized by that of ZO1. *p<0.05, unpaired t-test. (B) PK1 knockdown does not affect the ZO1 expression level. ns, not significant, unpaired t-test. (C) PK1 knockdown results in sustained defects in posterior neuropore (PNP) closure at 17ss.

**Figure S2.**
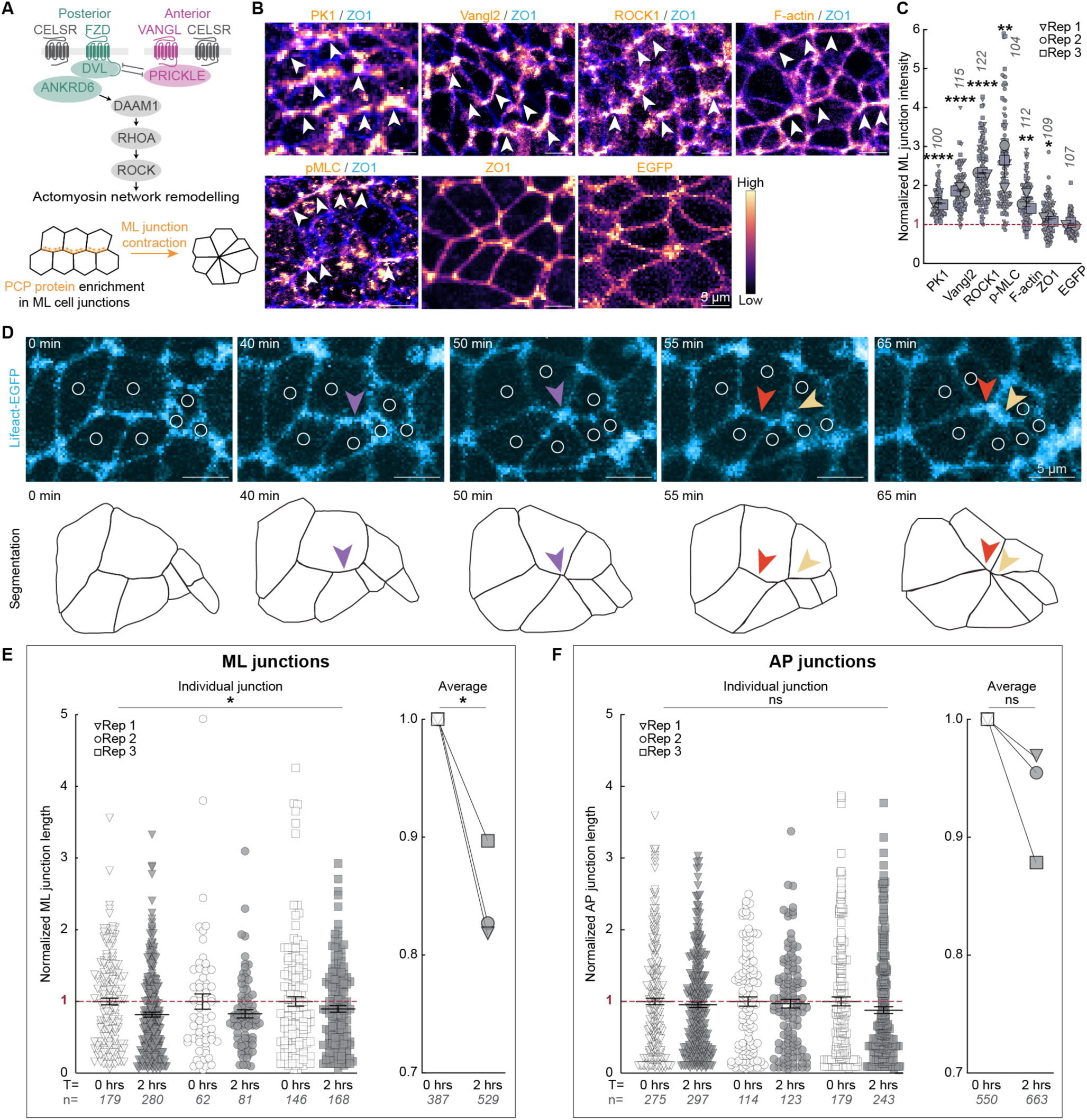
PCP signalling components are polarized along mediolateral cell junctions, related to Figure 2. (A) Schematic of Planar Cell Polarity signalling for actomyosin remodeling. (B, C) PCP components are preferentially enriched along the mediolateral (ML) cell junctions. The polarity of PCP proteins is compared to that of the membrane EGFP from transgenic quail embryos, which exhibits no apparent subcellular polarisation. ****p<0.0001, **p<0.01, *p<0.05, unpaired t-test. (D-F) The ML but not the anteroposterior (AP) cell junctions shrink in the JZ. The constricting ML cellular junctions are indicated by arrowheads in D. *p<0.05, ns, not significant, paired t-test.

**Figure S3.**
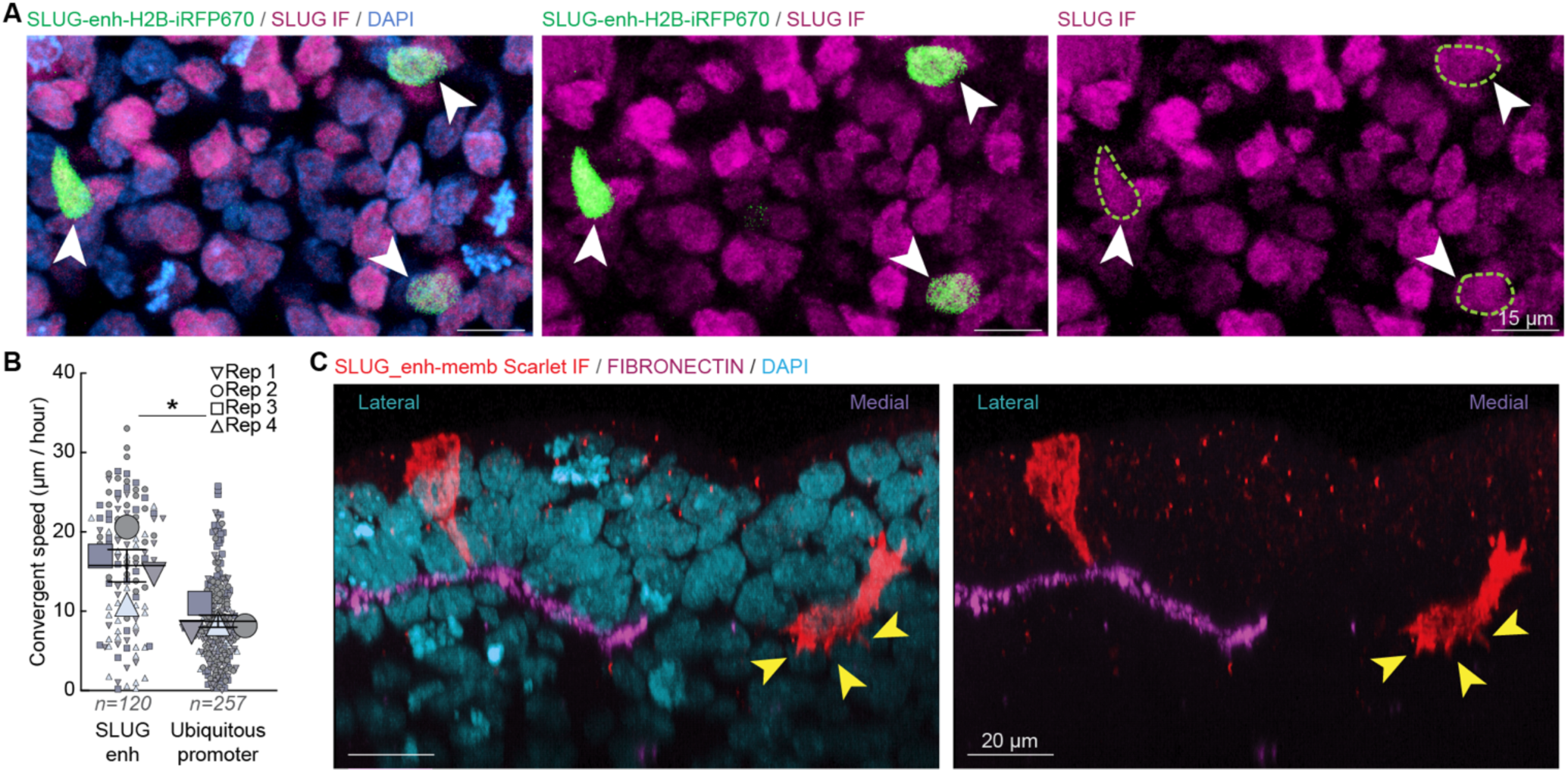
SLUG enhancer labels SLUG^+^ cells, related to Figure 4. (A) Cells expressing H2B-iRFP670 under the control of the SLUG enhancer (SLUG enh-H2B-iRFP670) are also positive for SLUG Immunofluorescent staining. (B) SLUG enhancer-positive cells converge significantly faster than cells labelled with ubiquitous CMV promoter. *p<0.05, unpaired t-test. (C) SLUG enhancer-positive cells have an epithelial morphology in the lateral JZ but a protrusive mesenchymal morphology (arrowheads) in the medial JZ.

**Figure S4.**
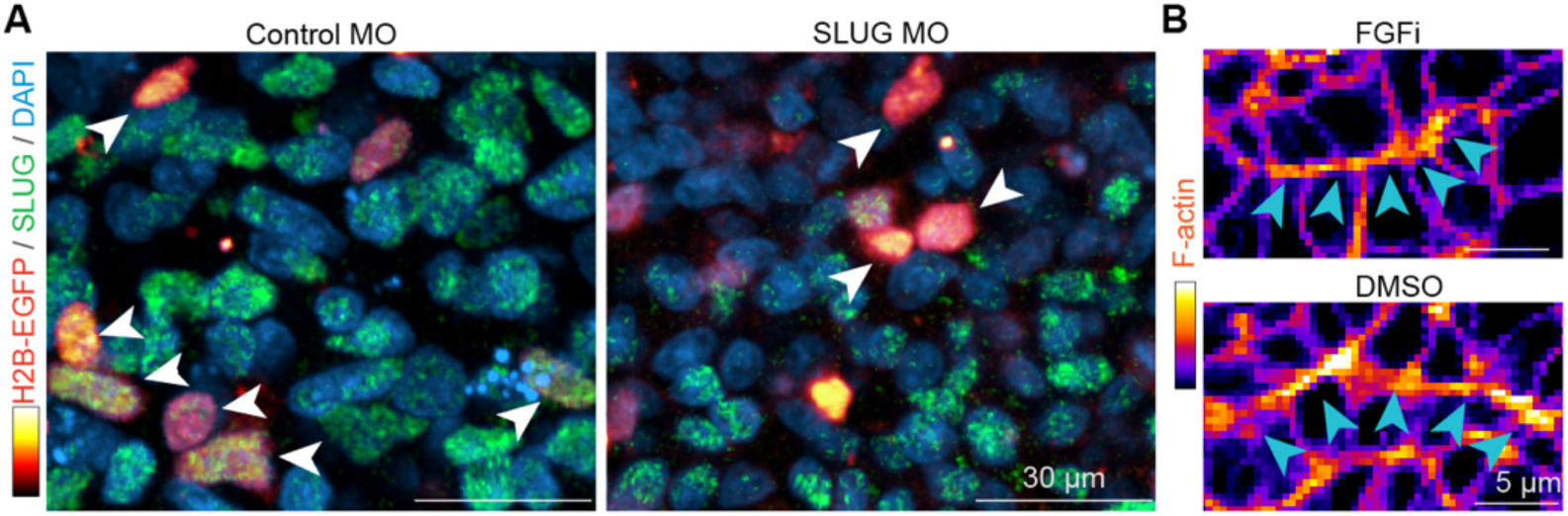
Controls for disrupting SLUG expression and FGF signalling, related to Figure 5. (A) Electroporation of a SLUG morpholino (MO) effectively reduces the number of SLUG+ cells in the JZ. The morpholino is co-electroporated with H2B-EGFP (in red). The arrowheads indicate cells double positive for SLUG staining and either a control MO or the SLUG MO. (B) FGF-inhibited embryos still assemble F-actin cables (arrowheads) in the lateral JZ.

**Figure S5.**
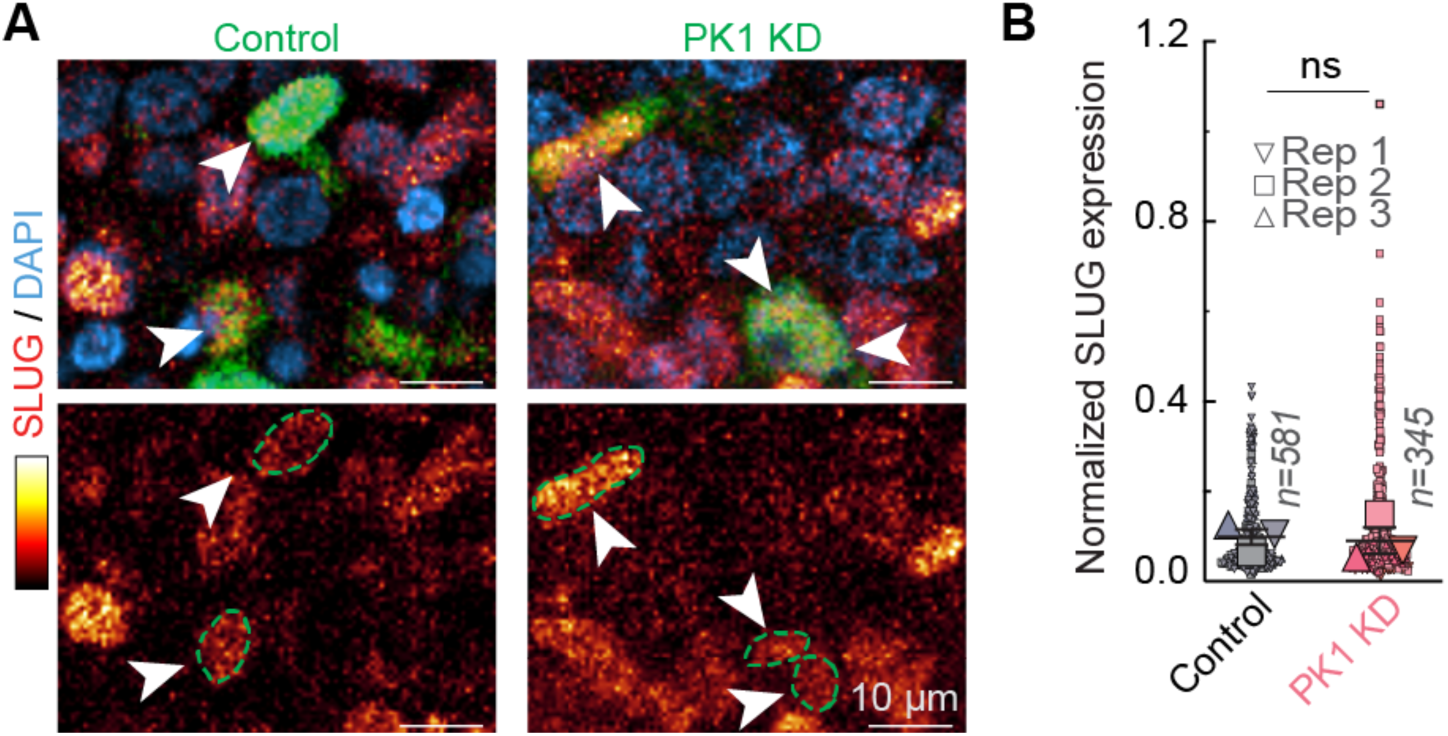
PK1 knockdown does not alter SLUG expression, related to Figure 6. (A) Immunofluorescent staining for SLUG reveals that Control and PK1 knockdown cells (green) still express SLUG (white arrowheads). (B) Quantification shows no change in SLUG expression following PK1 knockdown compared with Control (n = 3 embryos per group). ns, not significant, unpaired t-test.

**Figure S6.**
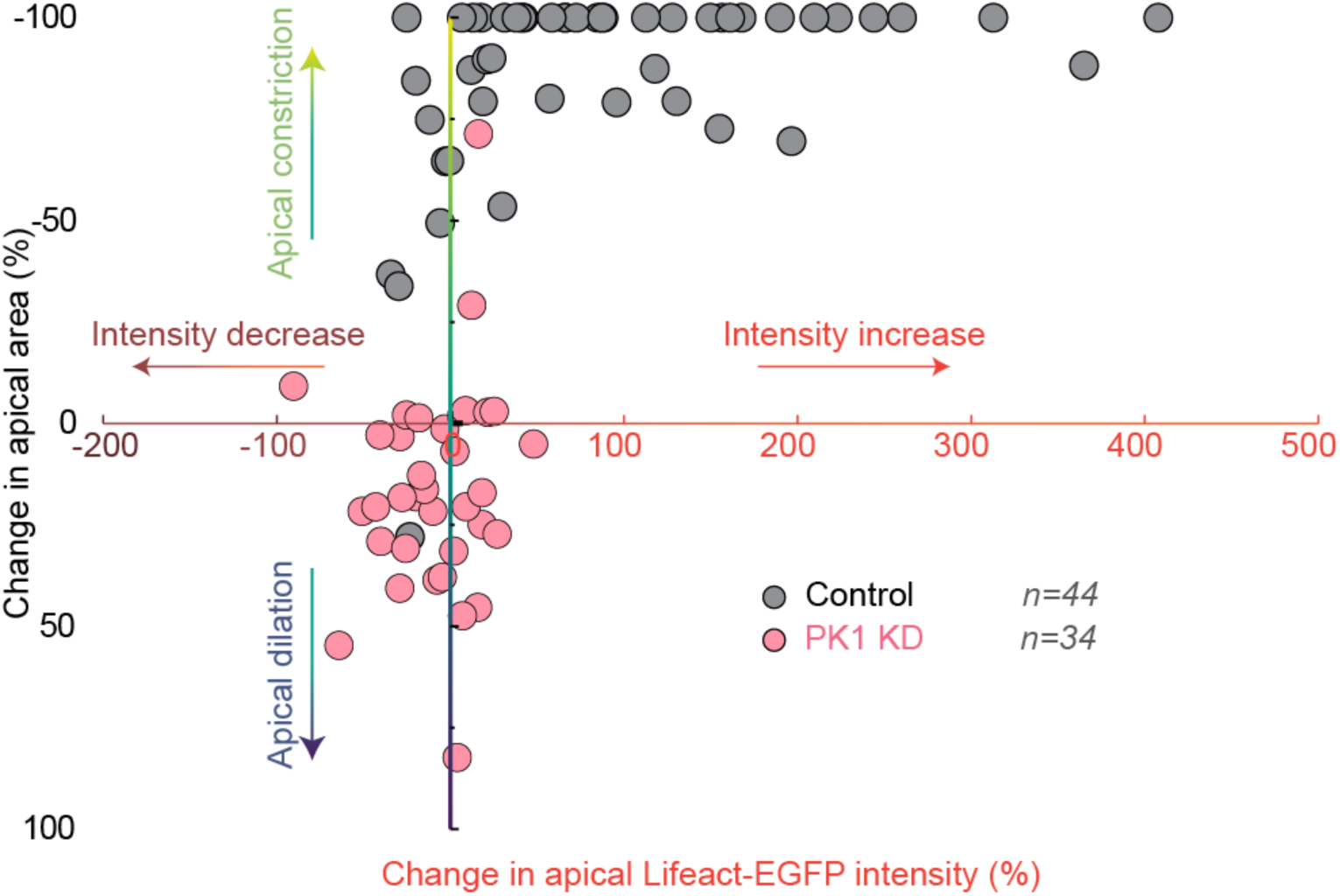
PK1 knockdown inhibits both the cortical F-actin enrichment and apical constriction, related to Figure 7. Live imaging of electroporated Lifeact-EGFP quail embryos reveals increased Lifeact-EGFP intensity at the apical cortex of Scrambled control cells as their apical surface constricts. PK1 knockdown cells fail to increase Lifeact-EGFP at their apical surface and do not constrict.

## SUPPLEMENTAL INFORMATION TITLES AND LEGENDS

**Video S1. JZ converges without extension, related to Figure 1**

A timelapse video of a Lifeact-EGFP quail embryo shows the JZ (outlined with white dotted line) across time. The triangle indicates the node. s, somite; nt, neural tube.

**Video S2. Cells in JZ converge, related to Figure 1**

This video shows individual tracks of migrating H2B-iRFP670 labelled cell nuclei in the frontal plane of the JZ. As the JZ is axisymmetric about the midline (white dotted line), the tracks are displayed on the right-hand side of the JZ for simplicity. The cell tracks are colour-coded by their medial displacement.

**Video S3. PK1 knockdown abolishes PNP closure, related to Figure 1**

Timelapse videos show PK1 knockdown results in junctional NTDs characterised by a wider posterior neuropore (PNP, outlined by white dotted line). The position of the last somite (S) is indicated by an asterisk.

**Video S4. PK1 knockdown disrupts cellular convergence, related to Figure 2**

Live imaging shows individual cell migration within the frontal plane in PRICKLE1 knockdown and control embryos. The cell tracks are colour-coded by their medial displacement. The triangle indicates the node and the white dotted line shows the midline.

**Video S5. Cells in medial JZ ingress, related to Figure 3**

This video shows individual tracks of migrating H2B-iRFP670 labelled cell nuclei in the transverse plane of the JZ. The cell tracks are colour-coded by their ventral displacement (ingression).

**Video S6. PK1 knockdown impairs medial cellular ingression, related to Figure 3**

Live imaging shows migration of individual cells within the transverse plane of PK1 knockdown and control embryos. The cell tracks are colour-coded by their ventral displacement (ingression).

**Video S7. SLUG positive cells migrate medially and ingress, related to Figure 4**

Live imaging shows SLUG positive cells (red) migrate towards the medial area of the JZ and ingress (green triangles).

**Video S8. PK1 knockdown perturbs cell apical constriction and ingression, related to Figure 6**

Timelapse videos show PK1 knockdown results in defects in cell apical constriction and ingression. The magenta triangles indicate the cells undergoing apical constriction and ingression.

## Notes

### Competing Interest Statement

The authors have declared no competing interest.

